# Evaluation validation of a qPCR curve analysis method and conventional approaches

**DOI:** 10.1101/2020.06.18.158873

**Authors:** Yashu Zhang, Hongping Li, Shucheng Shang, Shuoyu Meng, Ting Lin, Yanhui Zhang

**Author notes:** Hongping Li (HL).

## Abstract

Real-time quantitative polymerase chain reaction (qPCR) is a sensitive and reliable method for mRNA quantification and rapid analysis of gene expression from a large number of starting templates. It is based on the statistical significance of the beginning of exponential phase in real-time PCR kinetics, reflecting quantitative cycle of the initial target quantity and the efficiency of the PCR reaction (the fold increase of product per cycle). We used the large clinical biomaker dataset and 94-replicates-4-dilutions set which was published previously as research tools, then proposed a new qPCR curve analysis method——C_q_MAN, to determine the position of quantitative cycle as well as the efficiency of the PCR reaction and applied in the calculations. To verify algorithm performance, 20 genes from biomarker and partial data with concentration gradients from 94-replicates-4-dilutions set of MYCN gene were used to compare various publicly available curve analysis methods with our method and established a suitable evaluation index system. The results show that C_q_MAN method is comparable to other methods and can be a feasible method which applied to our self-developed qPCR data processing and analysis software, providing a simple tool for qPCR analysis.

## Introduction

The working principle of the qPCR is to add fluorophore into the qPCR system, and use the fluorescence signal accumulation to detect the whole qPCR process [1]. The accumulated amount of DNA reaction products after fluorescent labeling is used as amplification data (expressed as amplification curves) can be used to determine the initial target quantity (called N_0_ at the concentration level and called F_0_ at the fluorescence level). An amplification reaction is generally displayed by an amplification curve, while the y-axis represents the fluorescence signal accumulation and the x-axis represents the number of cycles. During the process, the product fluorescence can’t rise above the background at the beginning and almost tending to a straight line; as the reaction progresses, the amplification curve of the cumulative products show an S-typed increase [2-3]. The reason for this process is that, initially, the product quantity is very small, caused a weak fluorescence signal to be detected at baseline phase; then the products began to increase exponentially, and the amplification efficiency is closest to 100%. During the linear phase products continue to accumulate, but the reaction efficiency begins to fall and reagents become limiting. Until the product is no longer produced, so the reaction reaches to plateau phase [4]. Therefore, the baseline phase, exponential phase, linear phase, and plateau phase of the amplification curve are generated based on the quantitative relationship between the fluorescence signal accumulation and cycles in Fig 1.

**Fig 1.**
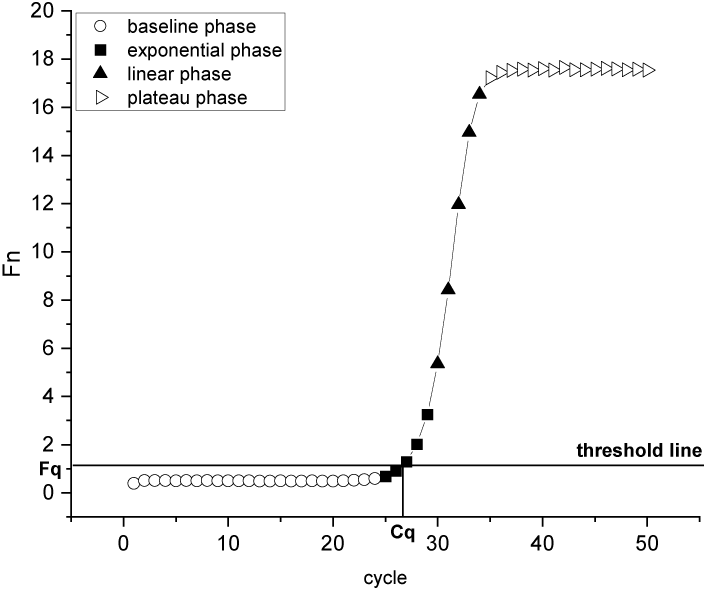
Four phases of the amplification curves. The initial fluorescence of the reaction is at the background level with high noise, almost no fluorescence signal can be detected, then the product fluorescence rises above the background in the exponential phase within a few cycles and begins to saturate in the approach to the final plateau phase.

For the relevant parameters of the amplification curve, the amplification process produces a quantitative threshold (called F_q_ in most methods) indicates a detectable fluorescent signal produced by the accumulation of sufficient amplification products. The x-axis of this quantitative threshold corresponds to a cycle called C_q_ in most methods, which is called C_q_MAN in our method.

The amplification efficiency(E) is another important parameter for checking qPCR data analysis. Under ideal conditions, the number of DNA sequences will double in each cycle, the percentage of E-1 is 100% (at this time E is 2) [5]. However, due to factors such as reaction inhibitors, enzyme, primer and probes differences, PCR efficiency rarely reaches 100%. Therefore, E is any number between 1 and 2[6]. Previously published studies have been suggested that PCR efficiencies mostly range between 65% and 90% [7].

After determining the quantitative cycle, the quantitative threshold, and estimating the amplification efficiency, the kinetics of qPCR exponential phase are described by Eq.1 to indicate the initial target quantity of the reaction.

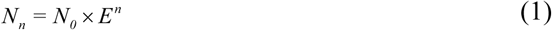

in which N_0_ and N_n_, are the initial target amount of DNA and the DNA target amount after n cycles, respectively. F_n_, the fluorescent signal after n cycles and F_0_, the fluorescence signal represents starting amount of the target DNA are the performance of N_n_ and N_0_ at the fluorescence level [2]. Therefore, Eq.1 can be described as Eq.2

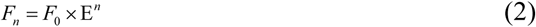

using the relevant parameters estimated by the curve analysis algorithm method can be expressed as Eq.3

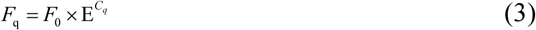

then the observed initial target quantity(F_0_) is calculated.

In the past two decades, the rapid development of qPCR technology has led to the production of multiple protocols, reagents, analytical methods and reporting formats [8]. However, most single amplification curve analysis methods that do not require the preparation of standard samples differ in determining the quantitative cycle (called C_q_ or C_t_ or other names and referred to C_q_MAN in our method) and estimating PCR reaction efficiency.In order to provide reference for further developing and evaluating the qPCR curve analysis method and promoting the research of quantitative fluorescence PCR in gene expression, the new curve analysis method and other methods were evaluated on the biomarker dataset and 94-replicates-4-dilutions set in this paper from the aspects of expression level and statistical significance. The goal of this paper is to make our new method comparable to other methods, at the same time provide users with an alternative curve analysis scheme. In order to evaluate the new method, some evaluation performance indicators were proposed.

## Materials and methods

### qPCR dataset

#### Biomaker dataset

Data comes from a previously published study that developed and validated the expression profile of a 59-mRNA gene to improve prognosis in children with neuroblastoma. This dataset measured 59 biomarkers and 5 reference genes in a sample maximization experimental design, using the LightCycler480 SYBR Green Master (Roche) in a 384-well plate with 8 μl reaction. These genes have been reported in at least two independent studies as prognostic genes for neuroblastoma. 366 cDNA samples from the primary tumor biopsy and a 5-point 10-fold serial dilution series based on an external oligonucleotide standards (from 150,000 to 15 copies, n = 3), and no template control (NTC, n = 3) are included in each plate [8]. This dataset will be referred to as ‘biomarker dataset’ in this study and 20 genes were selected from this dataset for analysis.

#### 94-replicates-4-dilutions set

This data set created a dilution series consisting of four 10-fold serial dilution points from 15,000 to 15 molecules, using 10 ng / µl yeast tRNA as a carrier (Roche) and created NTC samples of the same dilution. qPCR was done on a CFX 384 instrument (Bio-Rad). QPCR was performed on a CFX 384 instrument (Bio-Rad) using a 96-well pipetting robot (Tecan Freedom Evo 150). Amplification reactions were performed in 8µl samples containing) 0.4µl forward and 0.4µl reverse primer (5µM each), 0.2µl nuclease-free water, 4µl iQ SYBR Green Supermix (Bio-Rad) and 3µl of standard oligonucleotide. In 384-well plates (Hard-Shell 384-well microplate and Microseal B clear using an adhesive seal (Bio-Rad)), for each of the 4 dilution points, a total of 94 replicate reactions were distributed. In addition, the NTC reaction was repeated 8 times [9]. This dataset will be referred to as ‘94-replicates-4-dilutions set’. Since our system has 96 reaction channels, we select 44 curve data with concentration gradient from data set for analysis.

### qPCR curve analysis method

#### Previously published curve analysis method

The original standard-C_q_ method [10,11] fits a standard curve by preparing multiple sets of replicable experiments of the samples of known concentration, and estimates the concentration of unknown samples from the standard curve. Later, the exponential properties of the sigmoidal growth curve was determined to contain information on amplification efficiency. The efficiency can be calculated from the kinetics of a single PCR reaction and the initial target quantity of reaction can be estimated. Under this background, the LinRegPCR method [12] manually sets the linear window (W-o-L) to estimate the exponential phase; DART [13] constructs a model based on the maximum fluorescence value (R_max_) and the baseline fluorescence noise (R_noise_) to determine a central point M, and analyzes the cycle within a 10-fold range around M as an exponential phase; FPLM[14] uses a logistic approximation of the fluorescence curve to identify the exponential phase of a qPCR amplification plot; the bilinear model and the four-parameter logistic model are used in the FPK-PCR to estimate the initial target quantity [15]; 5PSM method [16] uses the ratio of the period of the first second derivative maximum (SDM) of the five-parameter S-type function to the previous period as the amplification efficiency; PCR-Miner[17] defines the exponential phase starting from 10-fold standard deviation of the baseline fluorescence noise and the first derivative maximum (SDM) of the four-parameter logistic model as the end;Cy0[18] obtains the intersection point between the abscissa axis of the curve inflection point and the tangent line based on the nonlinear regression of the Richards equation to the fluorescence value, and then estimates the initial target quantity.

#### C_q_MAN

C_q_MAN is an adaptive analysis system that summarizes the methods and experiences of previous methods and provides a robust, objective, and noise-resistant method for quantification of qPCR results that is independent of the specific equipment used to perform PCR reactions. Since researches have shown that smoothing can at best lead to erroneous accuracy of results, and usually also bias the results [20], the improved adaptive Savitzky-Golay filter in the C_q_MAN system is only used for visual display of data. The calculation of the method is based on the definition of the exponential phase of the amplification curve, reactions use the ‘midpoint’ of the exponential phase as the quantitative cycle. In the process of estimating single reaction E, the three-parameter single exponential fitting is performed from the “midpoint” to the end of the exponential phase defined as C_q_MAN. For multiple replicated experiments with concentration gradients,the estimate of E is obtained from the slope of the standard curve. At the same time, the average value of amplification efficiency (E_mean_) of the same gene is used when calculating the observed target quantity (F_0_) to ensure that the initial target quantity estimate is more reliable.

##### (1) Improved adaptive Savitzky-Golay filter

Savitzky-Golay is a filter based on the local polynomial least squares convolution algorithm in the time domain. It reduces noise while better maintaining the shape of the signal, and retains the distribution characteristics of the data points such as relative maximum, minimum, and width [20].

C_q_MAN system sets the width n of the filter window to 5 (n=2m+1), slide in steps of 5 and fits fluorescence values y_m_, y_-m+1_, …, y_0_, y_1_, …, y_m-1_, y_m_ within the window width, using k-1 degree polynomial(Eq.4) for fitting

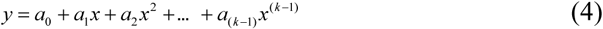

Therefore, there are n equations to constitute the k-ary linear in Eq.5. To make the equations have a solution, n should be greater than or equal to k, generally choose n>k, set to 4 in the system, and determine the fitting parameters a_0_, a_1_, …, a_k-1_ by least squares.

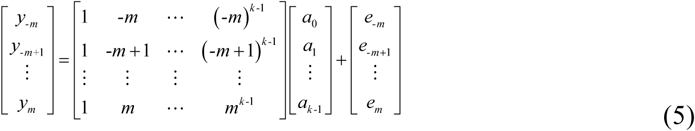

The matrix form of Eq. 5 is expressed as

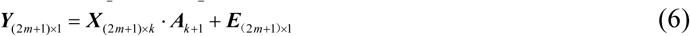

The least squares solution A’ of A is

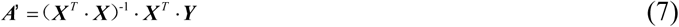

And the filtered fluorescence value matrix Y’ is

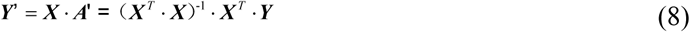

After that, the improved adaptive Savitzky-Golay filter makes the reconstructed amplification curve obtain the best filtering effect with the smallest fitting index(P_k_) through an iterative process.

a. Before the iteration, the first Savitzky-Golay smoothing process is performed to obtain the total trend line (Y^t^) of the curve.
b. The k-th iteration operation replaces the value of the trend line with a value lower than the total trend line to generate a new sequence (Y^k^).
c. Use the iteration result with the smallest fitting index (P_k)_ after the k-th iteration as the final filtering result.

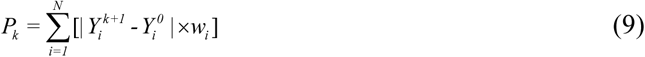

In Eq.9, 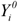 and 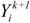 represent the i-th fluorescence value without iteration and the i-th fluorescence value after k-th iteration, w_i_ represents the weight of the i-th fluorescence value. The method for determining the weight is as follows:

a. The weight of all fluorescence values at the first smoothing is 1.
b. When iterating, w_i_ is obtained from Eq.10.

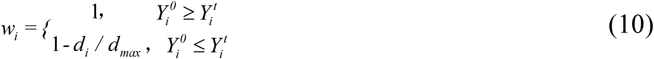

In Eq.10, 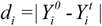 represents the distance between the uniterated i-th fluorescence value and the total trend line, and *d*_*max*_ represents the distance between the largest uniterated i-th fluorescence value and the total trend line.

##### (2) C_q_MAN method

C_q_MAN method relies on the four-parameter logistic model from the previous published method [14,17]. The four-parameter sigmoidal function is fitted to the raw fluorescence data by means of a non-linear fitting routine the Levenberg-Marquardt algorithm that minimizes the residual sum-of-squares to obtain parameters baseline fluorescence (y_0_), the difference between the maximum fluorescence and the baseline fluorescence (y_max_-y_0_), curve inflection point(x_0_) and b that affect the shape of amplification curve (Eq.11).

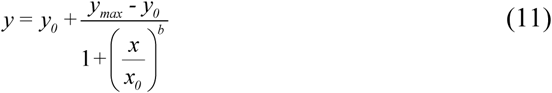

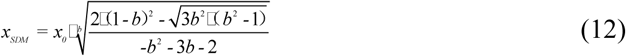

x_SDM_ is the point where the third derivative of the model is 0, that is, the cycle of second derivative maximum(SDM). The x_SDM_ cycle is applied as the end point of the exponential phase and the fluorescence value corresponding to this cycle is F_SDM_ in C_q_MAN method. Take the intermediate value of y_0_ and F_SDM_ as the quantitative threshold F_q_(Eq.13), then substitute this value into Eq. 11 to obtain the quantitative cycle (C_q_MAN).

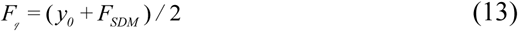

For efficiency estimation, a three-parameter simple exponent model is fitted to this exponential phase (from C_q_MAN to x_SDM_) using an iterative non-linear regression algorithm to estimate the single reaction’s individual efficiency in Eq.14, then the observed target quantity (F_0_) can be calculated by Eq.15.

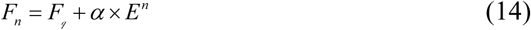

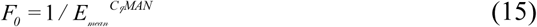

For multiple replicated experiments with concentration gradients, the C_q_MAN system uses least squares to construct a standard curve (Eq.16 as the logarithmic form of Eq.1) to estimate the efficiency E(Eq.17). Then the observed target quantity (F_0_) also can be calculated by Eq.15.

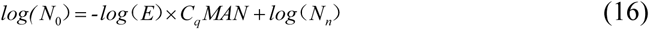

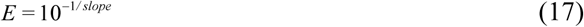

## Results and Discussion

### Performance Indicators

Performance analysis used 300 (5×3×20) amplification curve data of 20 genes with concentration of 150000, 15000, 1500, 150, 15 in biomaker and 3 replicated experiments for each group; 94-replicates-4-dilutions data set of 44 (4×11) amplification curves of the MYCN gene with a diluted concentration of 15, 150, 1500,15000,and each group of replicated experiments were 11 times. To eliminate the different measurement scales used by the analytical method based on concentration levels and fluorescence levels [21], we divided the data of all concentrations by the hightest concentration data and all fluorescence data by the average value of the maximum observed target quantity (F_0_), so that the average value of the maximum concentration and the maximum observed target quantity is 1. This process is called normalization. Then data sets were used to establish 4 performance indicators to measure the degree of compliance between the observed initial target quantity F_0_ and the true value calculated by the algorithm from different angles. Among them, the bias and relative error are used to compare the difference between the observed initial target quantity and the true value; coefficient of variation and precision are used to compare the difference between the observed initial target quantity of the same group. The smaller the difference, the more reliable the method. Performance indicators as follows.

#### (1) Bias

The ratio between the average of the observed initial target quantity F_0_ corresponding to the highest and lowest concentrations is calculated. In biomarker, the expected value of this ratio is 10,000 (because the ratio of the concentration of 150000 and 15 is 10000), and in 94-replicates-4-dilutions set, the expected value of this ratio is 0.001 (because the ratio of the diluted concentration of 15 and 15000 is 0.001) and any value deviating from 10,000 or 0.001 is expressed as a bias. The log-transformed (base 10) between the true value and the initial target quantity F_0_. After the data is normalized, the linear regression analysis makes the log (F_0_) and log(SQ) (SQ is the true concentration after normalization) slopes of the unbiased method 1 and any slope deviates from the value of 1 also expressed as a bias.

#### (2) Relative error (RE)

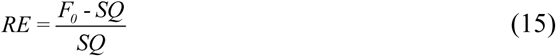

RE is the deviation after F_0_ and SQ are normalized to the same measurement scale.

#### (3) Coefficient of variation (CV)

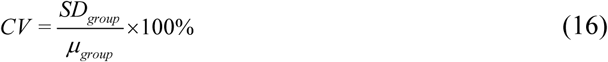

CV represents the ratio of the standard deviation (SD) to the average value(μ) of the same group (replicated experiments) of observed initial target quantity (F_0_).

#### (4) Precision

Precision represents the within-triplicate variance of the observed initial target quantity(F_0_) in the same group.

### Indicator Evaluation

#### Biomaker dataset analysis

##### (1) Bias

We expect the ratio between the observed initial target quantity and the true value to be 10,000 or 0.001 in two different datasets. After the data is normalized, the linear regression analysis makes the log (F_0_) and log (SQ) slopes of the unbiased method 1, which will be unbiased. C_q_MAN, Cy0 has an advantage in the deviation index because the two methods calculate the efficiency value based on the slope of the relationship between Cy0 and log (input), and then use this efficiency value and the Cy0 value to calculate F_0_. Therefore, this methods are unbiased and are the result of cyclical reasoning, but this also ensures that the observed initial target quantity F_0_ is more accurate. Other methods are positively or negatively biased, and the observed values deviate significantly from the true values.

##### (2) Relative error

The relative error was originally used to compare the difference between the measured value and the true value, and the degree of confidence in the response measurement. Here we can use the relative error response to calculate the difference between the observed value and the true value, reflecting the credibility of the algorithm. More intuitive response measurement accuracy than absolute error. We use relative error as one of the indicators to determine the difference between the observed initial target quantity F_0_ and the true value. Cy0 performed best, average relative error was 0.1050. The average relative error of the rank after the second PCR-Miner was 0.2287, C_q_MAN was 0.2416, and the highest 5PSM was as high as 0.6939.

##### (3) Coefficient of variation

The coefficient of variation reflects the degree of dispersion of the data, and at the same time overcomes the effects of large differences in measurement scales or different data sizes. We use the coefficient of variation coefficient to calculate the degree of dispersion of the observed initial target quantities of the three groups at each concentration, and average the five groups of coefficients of variation. The smaller the coefficient of variation, the lower the degree of dispersion. Result showed that C_q_MAN showed the best performance of 7.20%, Cy0, LinRegPCR, PCR-Miner also stabilized at about 9.60%, and FPK-PCR’s coefficient of variation was as high as 25.12% in Table 1.

**Table 1.**
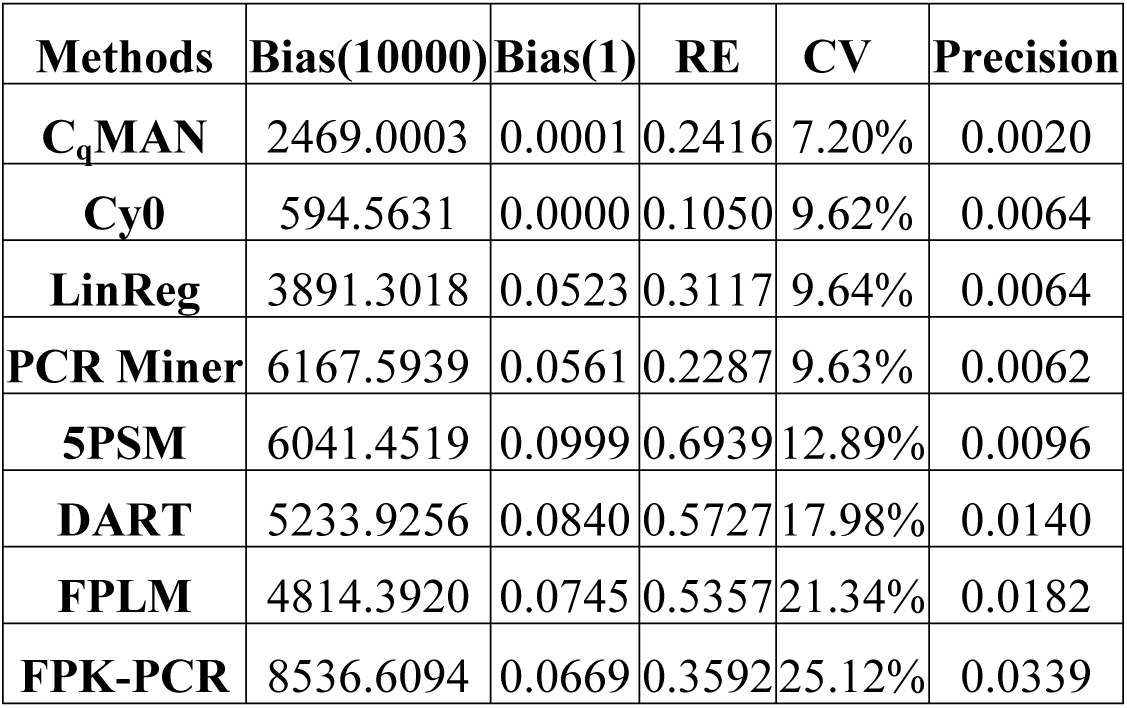
Analysis of the average of 20 genes in 4 indicators per method.

| Methods | Bias(10000) | Bias(1) | RE | CV | Precision |
| --- | --- | --- | --- | --- | --- |
| **C<sub>q</sub>MAN** | 2469.0003 | 0.0001 | 0.2416 | 7.20% | 0.0020 |
| **Cy0** | 594.5631 | 0.0000 | 0.1050 | 9.62% | 0.0064 |
| **LinReg** | 3891.3018 | 0.0523 | 0.3117 | 9.64% | 0.0064 |
| **PCR Miner** | 6167.5939 | 0.0561 | 0.2287 | 9.63% | 0.0062 |
| **5PSM** | 6041.4519 | 0.0999 | 0.6939 | 12.89% | 0.0096 |
| **DART** | 5233.9256 | 0.0840 | 0.5727 | 17.98% | 0.0140 |
| **FPLM** | 4814.3920 | 0.0745 | 0.5357 | 21.34% | 0.0182 |
| **FPK-PCR** | 8536.6094 | 0.0669 | 0.3592 | 25.12% | 0.0339 |

##### (4) Precision

The five concentration sequences were measured three times and the fluorescence data were analyzed. Therefore, the variance of each set of 3 measurements should be small, reflecting only random changes in laboratory procedures and fluorescence measurements, and such changes should always be the same. The resulting three internal variances can be considered as a measure of the accuracy of the analytical method. C_q_MAN, 5PSM, Cy0, LinRegPCR have lower variability.

##### (5) Efficiency

The range of differences in efficiency values for each method indicates that this variability is the sum of the difference in efficiency between genes and the difference in estimation methods. Therefore, the difference between the methods cannot be explained. Except that DART and FPLM share a method of finding E, other methods get different median values of E. FPK-PCR has a large number of efficiency values above 2, which is obviously too high. The median value of PCR Miner is 1.11, which is significantly lower. The median value of C_q_MAN, Cy0, LinRegPCR, 5PSM is between 1.7 and 1.9. We calculated the standard deviation of the amplification efficiency of the 20 genes, in which LinRegPCR, DART, FPLM calculated E value is relatively stable in Fig 3.

**Fig 2.**
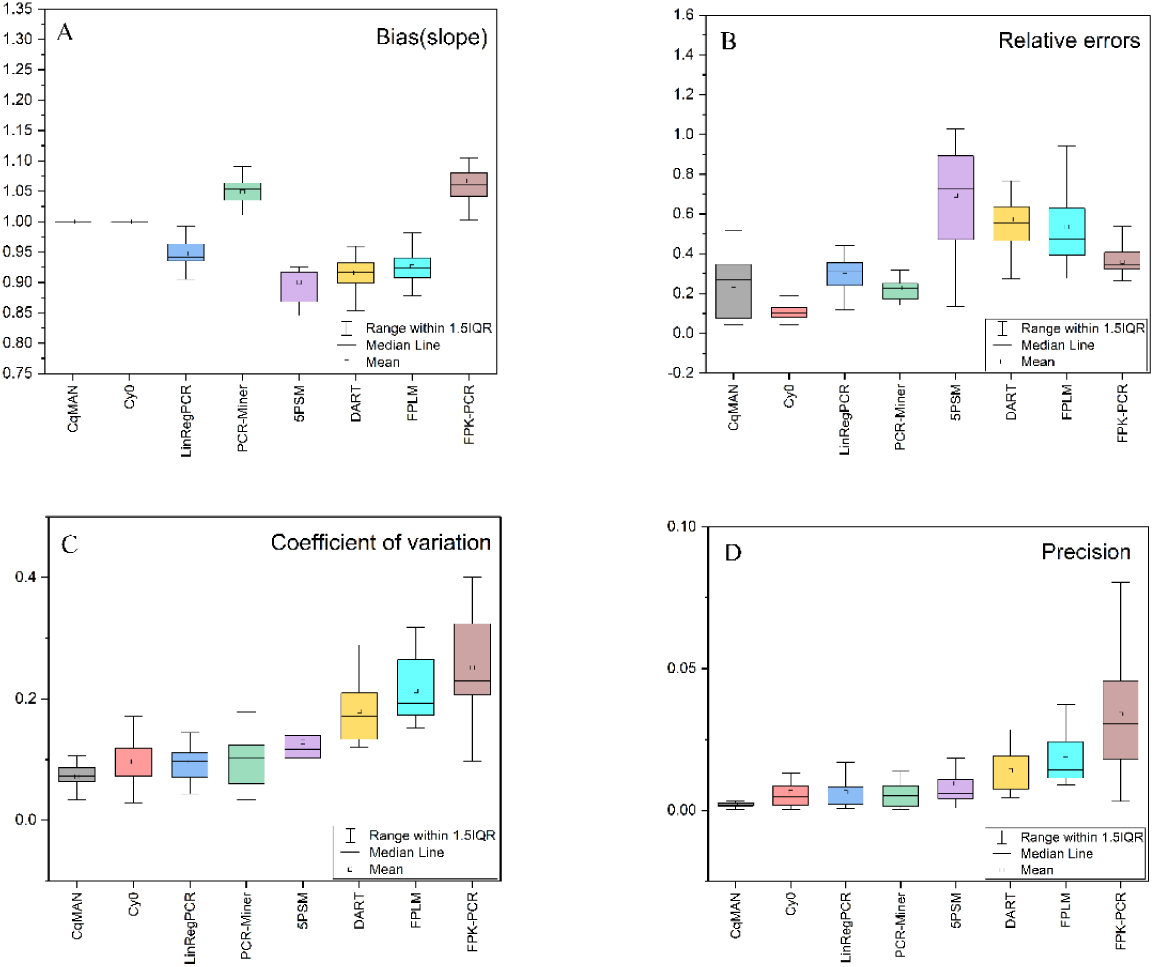
Performance indicators per method. The performance indicator values determined from the concentration series included in the measurement of the 20 genes are summarized in box-and-whisker plots. The boxes range from the 25th to the 75th percentile and are divided by the median; the whiskers are set at the 5th and 95th percentile (A)Bias in the slope level, which is based on the degree of deviation from 1.(B)The box-and-whisker plot of relative errors shows the difference between the observed initial target quantity and the true value.(C)Coefficient of variation is an objective indicator of the effects of measurement scales and dimensions that eliminate fluorescence levels and concentration levels.(D)Precision is determined as the within-triplicate variance and should have the same, low, value in all methods

**Fig 3.**
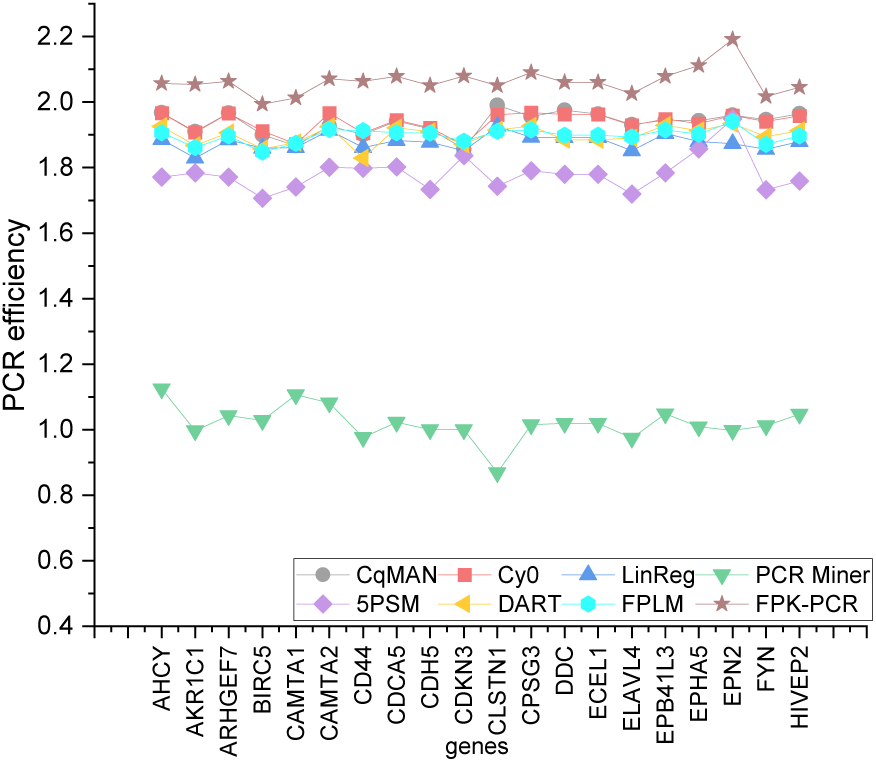
PCR efficiency per gene. The mean value of the efficiencies of each gene in per method.

#### 94-replicates-4-dilutions set analysis

##### (1) Target quantity

For data with dilute concentrations of 15000, 1500, 150, and 15, respectively, the observed target quantity should be as close as possible to the expected value −3, −2, −1, 0 obtained after calculating the log (F_0_) (base 10) in Fig 4. The systematic negative or positive deviation of each analysis method is shown by the deviation of the average F_0_ from the expected value (Fig 4: horizontal line). C_q_MAN, Cy0, PCR-Miner and LinRegPCR have the least bias. DART and FPLM show a higher bias, 5PSM displays a strong overestimation whereas FPK-PCR shows a strong underestimation of F_0_ values.

**Fig 4.**
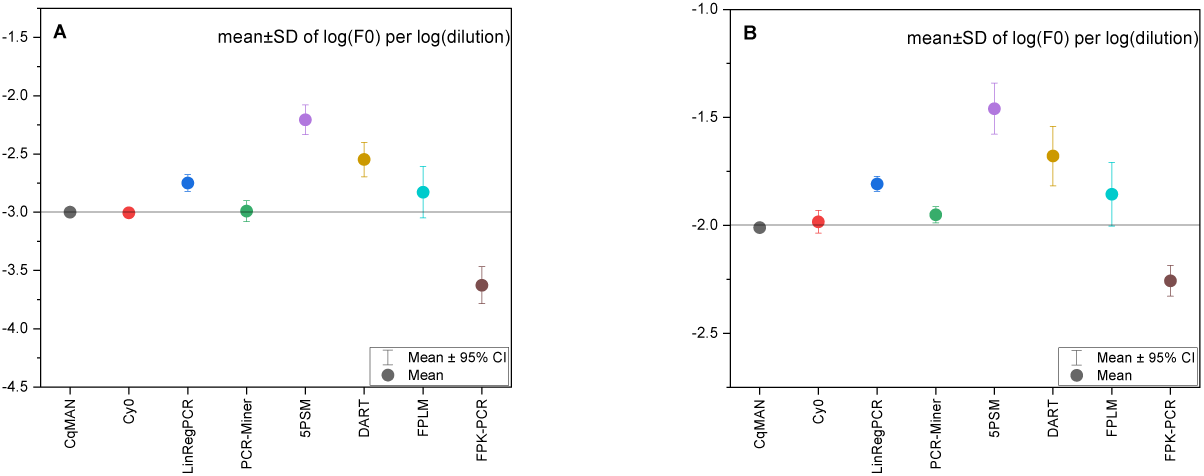
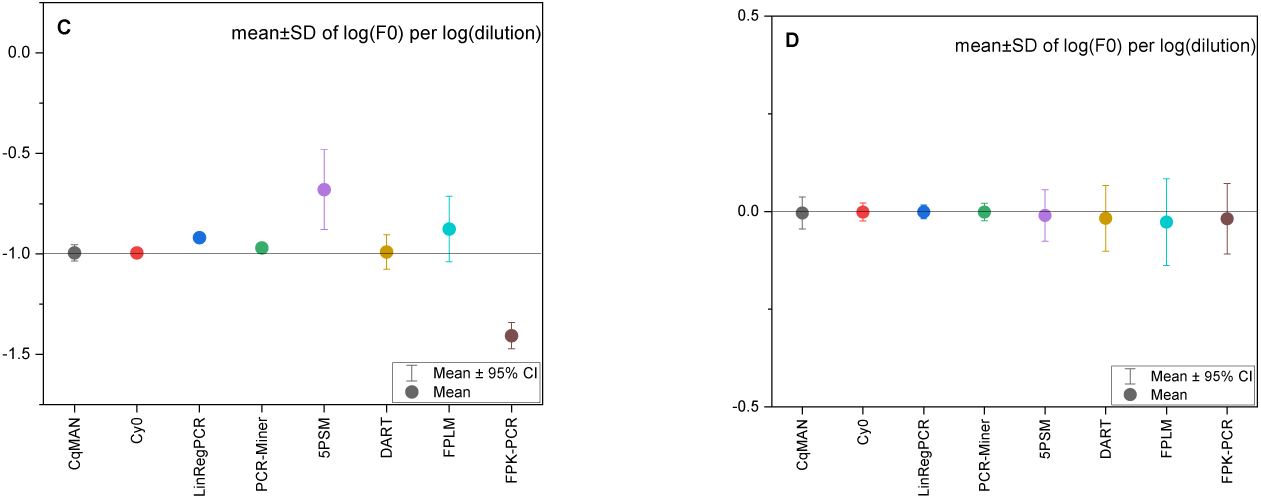
Mean observed F_0_ value per concentration and method. The highest dilute concentration is set to 1, the y-axis is set to log (dilution) (base 10).

(2)Bias, RE, CV, precision and E. C_q_MAN and Cy0 keep lower variance in bias. C_q_MAN perform best in RE, CV and precision. C_q_MAN, Cy0, LinRegPCR and PCR-Miner does not vary much between the values in CV and precision. Table 2 clearly illustrate the differences in 5 indicators of 8 methods and the average PCR efficiency of these methods is provided. The efficiency of Cy0 was not provided in the previously published data analysis.

**Table 2.** Analysis of MYCN gene in 4 indicators and the mean of PCR efficiency per method.

| Methods | Bias(0.001) | Bias(1) | RE | CV | Precision | $\bar{E}$ |
| --- | --- | --- | --- | --- | --- | --- |
| **C<sub>q</sub>MAN** | 0.0000 | 0.0000 | 0.0873 | 10.86% | 0.0025 | 1.7400 |
| **Cy0** | 0.0000 | 0.0000 | 0.0935 | 10.42% | 0.0027 | NA |
| **LinRegPCR** | 0.0008 | 0.0869 | 0.4126 | 11.32% | 0.0039 | 1.8690 |
| **PCR-Miner** | 0.0001 | 0.0054 | 0.1434 | 12.39% | 0.0057 | 1.9905 |
| **5PSM** | 0.0057 | 0.2633 | 2.5536 | 41.40% | 0.0409 | 1.7462 |
| **DART** | 0.0022 | 0.1723 | 1.0169 | 39.86% | 0.0308 | 1.9047 |
| **FPLM** | 0.0008 | 0.0617 | 0.6724 | 50.71% | 0.0609 | 1.9844 |
| **FPK-PCR** | 0.0007 | 0.1672 | 0.5012 | 29.55% | 0.0236 | 2.3011 |

For each of the evaluation indexes of the concentration sequence analysis of each gene, the rank synthesis method was used, and the Friedman test determined that these methods were not significantly different and comparable. Table 3 shows the results of each gene and method. The lower average rank indicates that the method which estimates the initial target quantity is closer to the true value in the performance evaluation of the four indicators we selected.

**Table 3.** Analysis of performance parameters per method in biomarker dataset (left) and 94-replicates-4-dilutions set(right). For each method, the mean rank is given for each of the performance indicators bias, RE, CV, precision. The methods are sorted based on the average of these ranks.

| **Methods** | **Bias(10000/0.001)** | **Bias(1)** | **RE** | **CV** | **Precision** | **rank** |
| --- | --- | --- | --- | --- | --- | --- |
| **$C_q$MAN** | 2/2 | 1/1 | 3/1 | 1/2 | 1/1 | 1.70/1.50 |
| **Cy0** | 1/1 | 1/1 | 1/2 | 2/1 | 4/2 | 1.90/1.50 |
| **LinRegPCR** | 3/6 | 2/4 | 4/4 | 4/3 | 3/3 | 3.40/4.20 |
| **PCR-Miner** | 7/3 | 3/2 | 2/3 | 3/4 | 2/4 | 3.60/3.40 |
| **5PSM** | 6/8 | 7/7 | 8/8 | 5/7 | 5/7 | 6.40/7.60 |
| **DART** | 5/7 | 6/6 | 7/7 | 6/6 | 6/6 | 6.20/6.60 |
| **FPLM** | 4/5 | 5/3 | 6/6 | 7/8 | 7/8 | 6.00/6.20 |
| **FPK-PCR** | 8/4 | 4/5 | 5/5 | 8/5 | 8/5 | 6.80/5.00 |

In the average rank sorting of 20 genes in the biomaker data set, the performance of C_q_MAN is 1.70, the average rank of Cy0 is 1.90. The rank averages of the 5PSM, DART, FPLM, and FPK-PCR are all above 6, and the overall performance of F_0_ estimation is lower in Table 3. For the 94-replicates-4-dilutions set, the lowest rank average of C_q_MAN and Cy0 are 1.50, and the performance of LinRegPCR and PCR-Miner is also good; the rank average of 5PSM, DART, FPLM, and FPK-PCR is much higher.

## Conclusions

Based on real-time PCR kinetics and exponential model simulations, this study summarizes the real-time quantitative PCR curve analysis method proposed by the predecessors, and proposes a reliable gene expression level quantification method, C_q_MAN. To prove the reliability of the method, two data sets from different instruments, different PCR mixtures, and a testable hypothesis were used to evaluate the performance of multiple qPCR curve analysis methods. The fluorescence data of the other 7 methods in the performance analysis process were taken from a previously published research by Ruijter et al. in 2013[9]. Since the supplemental information from this research provided an excel template for calculating bias and precison, we can directly import the amplification curve data from two data sets analyzed by the C_q_MAN system into the excel template to obtain the calculated values of the two indicators. The relative error and coefficient of variation are the two statistical indicators proposed by the author for evaluation and analysis. Therefore, due to the difference in indicator settings and the difference in data sets selection, our analysis results are different from the results previously published by Ruijter et.al.

The limitation of this study is that two datasets have limited evaluation of the general applicability of the C_q_MAN method, so future researches should include more instances and more verification indicators to better verify the robustness and representativeness of the method. However, it is undeniable that the analysis templates, datasets, and analysis results (see supporting information) in this research will definitely help further evaluation of research and make the results comparable with our results.

The aim of this study is not to promote a particular curve analysis method with the best overall performance, because the choice of methods by the experimenters may depend on the different research goals of experimental instruments, reagents, protocols, etc. It is our intention to help users choose the ideal method for their own studies and developers to modify and improve their methods.

Finally we provide the URL of the C_q_MAN system: http://122.193.29.190:9999/xMAN/en-us/index. All data sets and analysis conclusions described in the article are provided in supporting information.

## Acknowledgments

This work is supported by Apexbio Biotechnology(Suzhou) Co., Ltd. The authors wish to thank Professor Ting Lin’s group for their kindly assistance in providing the technical support.

## Supporting information

### S1 File. Biomaker_set

This file provides the raw fluorescence data of 20 genes in the biomaker dataset used to compare the performance of each method in the article. Each .xlsx file has been processed into a readable format of the C_q_MAN system.

### S2 File. 94_replicate_4_dilutions_set

This file provides the raw fluorescence data of MYCN in the 94_replicate_4_dilutions_set used to compare the performance of each method in the article. Each .xlsx file has been processed into a readable format of the C_q_MAN system.

### S3 File. Biomaker_set_results

This folder contains two .xlsx files. The biomaker_analysis_dilutoin_series contains three sheets that provide the calculation process of performance indicators (take C_q_MAN data as an example), F_0_ and C_q_ values of 8 methods. The biomaker_performance_indicators integrates the analysis results of 20 genes performance indicators.

### S4 File. 94_replicate_4_dilutions_set_results

This folder contains two .xlsx files. The 94_replicates_4_dilutions_set_performance indicators integrate the analysis results of performance indicators. The 94_replicates_4_dilutions_set_results provides the relevant parameters calculated by 8 methods.

## References

1. A. Tichopad, M. Dilger, G. Schwarz, and M.W. Pfaffl. Standardized determination of real-time PCR efficiency from a single reaction set-up. Nucleic Acids Res. 1993;31(20): e122.

2. R. Higuchi, C. Fockler, G. Dollinger, R. Watson. Kinetic PCR analysis: realtime monitoring of DNA amplifification reactions. Biotechnology. 1993;11: 1026–1030.

3. Joel Tellinghuisen, Andrej-Nikolai Spiess. Comparing real-time quantitative polymerase chain reaction analysis methods for precision, linearity, and accuracy of estimating amplification efficiency. Analytical Biochemistry. 2014;449:76–82.

4. Heather D. VanGuilder, Kent E. Vrana,Willard M. Freeman. Twenty-five years of quantitative PCR for gene expression analysis. BioTechniques. 2008;44(5):619–629.

5. Xiayu Rao, Dejian Lai, Xuelin Huang. A new method for quantitative real-time polymerase chain reaction data analysis. Journal of computational biology. 2013;20(9):703–711.

6. Liu, W., and Saint, D.A. A new quantitative method of real time reverse transcription polymerase chain reaction assay based on simulation of polymerase chain reaction kinetics. Anal. Biochem. 2002;302:52–59.

7. Kamphuis, W., Schneemann, A., van Beek, L.M., et al. Prostanoid receptor gene expression profile in human trabecular meshwork: a quantitative real-time PCR approach. Invest. Ophthalmol. Vis. Sci. 2001;42: 3209–3215.

8. J. Vermeulen, K. DePreter, A. Naranjo, L. Vercruysse, R.N. Van, J. Hellemans, K. Swerts, S. Bravo, P. Scaruffi, G.P. Tonini, B.B. De, R. Noguera, M. Piqueras, A. Canete, V. Castel, I. Janoueix-Lerosey, O. Delattre, G. Schleiermacher, J. Michon, V. Combaret, M. Fischer, A. Oberthuer, P.F. Ambros, K. Beiske, J. Benard, B. Marques, H. Rubie, J. Kohler, U. Potschger, R. Ladenstein, M.D. Hogarty, P. McGrady, W.B. London, G. Laureys, F. Speleman, J. Vandesompele. Predicting outcomes for children with neuroblastoma using a multigene-expression signature: a retrospective SIOPEN/COG/GPOH study. Lancet Oncol. 2009:10(7):663–671.

9. J.M. Ruijter, M.W. Pfaffl, S. Zhao, A.N. Spiess, G. Boggy, J. Blom, R.G. Rutledge, D. Sisti, A. Lievens, K. De Preter, S. Derveaux, J. Hellemans, J. Vandesompele. Evaluation of qPCR curve analysis methods for reliable biomarker discovery: Bias, resolution, precision, and implications. Methods. 2013;59(1):32–46.

10. Larionov, A., Krause, A., and Miller. A standard curve based method for relative real time PCR data processing. BMC Bioinformatics. 2005;6: e62.

11. J.M. Ruijter, C. Ramakers, W.M. Hoogaars, Y. Karlen, O. Bakker. LinRegPCR: Analysis of quantitative RT-PCR data [computer program]. Version 11.0. Amsterdam, the Netherlands: Heart failure research center. Academic Medical Centre,2003;37: e45.

12. J.M. Ruijter, C. Ramakers, W.M. Hoogaars, Y. Karlen, O. Bakker, M.J. van den Hoff, A.F. Moorman. Amplifification effificiency: linking baseline and bias in the analysis of quantitative PCR data. Nucleic Acids Research. 2009;37:e45.

13. Peirson Stuart N, Butler Jason N, Foster Russell G. Experimental validation of novel and conventional approaches to quantitative real-time PCR data analysis. Nucleic Acids Research. 2003;31(14): e45.

14. C.A. Heid, J. Stevens, K.J. Livak, et al,. Real time quantitative PCR. Genome Res. 1996;6: 986–994.

15. A. Lievens, A.S. Van, M. Van den Bulcke, E. Goetghebeur. Enhanced analysis of real-time PCR data by using a variable efficiency model: FPK-PCR. Nucleic Acids Research. 2012;40(2): e10.

16. C. Ritz, A.N. Spiess. qPCR: an R package for sigmoidal model selection in quantitative real-time polymerase chain reaction analysis. Bioinformaticas. 2008;24(13):1549–1551.

17. Spiess A N, Feig C,Ritz C. Hightly accurate sigmoidal fitting of real-time PCR data by introducing a parameter for asymmetry. BMC Bioinformatics. 2008;9: e211.

18. S. Zhao, R.D.Fernald, J. Comprehensive algorithm for quantitative real-time polymerase chain reaction. Journal of Computational Biology. 2005;12(8):1047–1064.

19. Guescini M., Sisti D., Rocchi M.B., et al. A new real-time PCR method to overcome significant quantitative inaccuracy due to slight amplification inhibition. BMC Bioinformatics. 2008;9: e326.

20. A.N. Spiess, C. Deutschmann, M. Burdukiewicz, R. Himmelreich, K. Klat, P. Schierack, S. Rödiger. Impact of smoothing on parameter estimation in quantitative DNA amplification experiments. Clinical Chemistry. 2015;61(2):379–388.

21. M. Vynck, O. Thas. Reducing bias in digital PCR quantification experiments: The importance of appropriate modelling of volume variability. Analytical Chemistry. 2018;90(11):6540–6547.

